# Paternal care as a source of key skin microbiota in a poison frog

**DOI:** 10.64898/2025.12.30.696907

**Authors:** Francesca N. Angiolani-Larrea, Rayan Bouchali, Adeline Loyau, Dirk S. Schmeller, Eva Ringler

## Abstract

Parental care is a well-established mechanism for the vertical transfer of gut microbiota, yet its role in shaping offspring skin microbial communities remains largely unexplored. Using a cross-fostering experiment in a Neotropical poison frog, we demonstrate that paternal tadpole transport is a critical vector for the transmission of skin microbiota. Tadpoles transported by both biological and foster fathers acquired microbial communities that persisted through metamorphosis, maintaining a stable membership despite shifts in relative abundance. Notably, the paternal contribution included multiple bacterial genera known to inhibit the lethal pathogen *Batrachochytrium dendrobatidis* (Bd). In contrast, tadpoles deprived of paternal transport and manually transferred to water bodies exhibited depauperate communities with a significantly lower prevalence of these protective bacteria. Our findings reveal that parental care is a multifaceted behaviour that not only ensures offspring survival but also actively provisions them with a tailored, protective skin microbiome, highlighting a novel mechanism by which parental care influences offspring fitness and resilience to disease.

## Introduction

Social behaviour was recently hypothesized to play a key role in transferring all kinds of materials between individual animals [1]. For example, during mating, grooming, or parental care, social partners or parents and offspring stay in close spatial association with each other, which might facilitate the transfer of chemicals, proteins, nutrients, as well as microbiota. Most organisms live and cooperate closely with huge numbers of symbiotic microorganisms which live both inside and outside of their body [2–4]. These microbiomes have been shown to hold key roles in the host health, as they stimulate or be a direct part of the immune system [5–8]. However, we have limited knowledge about the role of social interactions contributing to the building and development of the microbiome, especially in species with complex life cycles (cf. [9–11]). In humans, it has been shown that gut microbiota are acquired during delivery in the birth canal and through the mother’s breast milk [4; 12; 13]. In other animals, likewise, microbial transfers between parents and offspring have mainly been found in species where parents feed their young, such as mucus-feeding fish [14], pigeons that feed on crop milk [15], or skin-feeding caecilians [16], and also in social insects [17–19].

The skin holds an essential role for individual health, as it acts as a physical protective barrier against external threats such as pollutants or pathogens, and is associated with diverse microorganisms [3]. In amphibians, the skin is of particular importance as it regulates key body functions such as respiration, hydration, and immune responses [20–22]. Diversity of the amphibian skin microbiome leads to increased protection from lethal fungal pathogens, such as *Batrachochytrium dendrobatidis*, *Batrachochytrium salamandrivorans* [23–25], or ranaviruses [26], through metabolites produced by bacteria [20], or direct competition between the pathogen and bacteria [27]. Only little empirical evidence suggests a direct transfer of microbiota on the amphibian skin via close body contact during parental care (but see [28–31]). Moreover, many amphibian taxa undergo dramatic morphological and physiological changes during metamorphosis [32], which might constrain the effectiveness of such initial seeding over longer timescales. It is also not known if the composition and diversity of the acquired microbiota at earlier developmental stages is maintained beyond metamorphosis.

Amphibians hold an immense diversity of reproductive modes [33; 34], which offers a unique opportunity to study the role of social behaviours for microbiota acquisition. In Neotropical poison frogs, parental care is obligate for offspring survival, as clutches are deposited in the leaf litter or on other terrestrial structures, and the hatched tadpoles are then transported on the back of the father to water bodies where their development is completed [35–38]. Tadpole transport has long been viewed primarily as a behavioural adaptation to assure that terrestrial eggs develop into aquatic larvae [34]. During tadpole transport, prolonged skin-to-skin contact between fathers and offspring may facilitate skin microbiome transmission. A previous study found that substantial bacterial colonization of embryos takes place directly after hatching, and that tadpole transport indeed serves as a source of skin microbes for transported tadpoles [31]. However, this study did not find a higher degree of similarity between microbial communities of tadpoles and adults in species that transport their offspring compared to those that did not. Communities of tadpoles also didn’t show a higher similarity to their caregiver than to unrelated adults, indicating that most caregiver-associated microbes do not remain in tadpole communities long-term.

To test if parental care is a key source for the acquisition of beneficial skin microbiota, and if this initial seeding is maintained over metamorphosis, we used the Neotropical poison frog *Allobates femoralis*. We conducted a cross-fostering experiment under controlled laboratory conditions, where hatched tadpoles were transported either by their own father, by foster fathers, or manually transferred to the provided water bodies, to disentangle microbial acquisition from adult frogs and the environment (Fig. 1). All tadpoles were reared until metamorphosis to identify if acquired microbiota are indeed retained until this developmental stage, and to reveal potential benefits to microbial composition (e.g. persistence of protective microbes).

**Figure 1.**
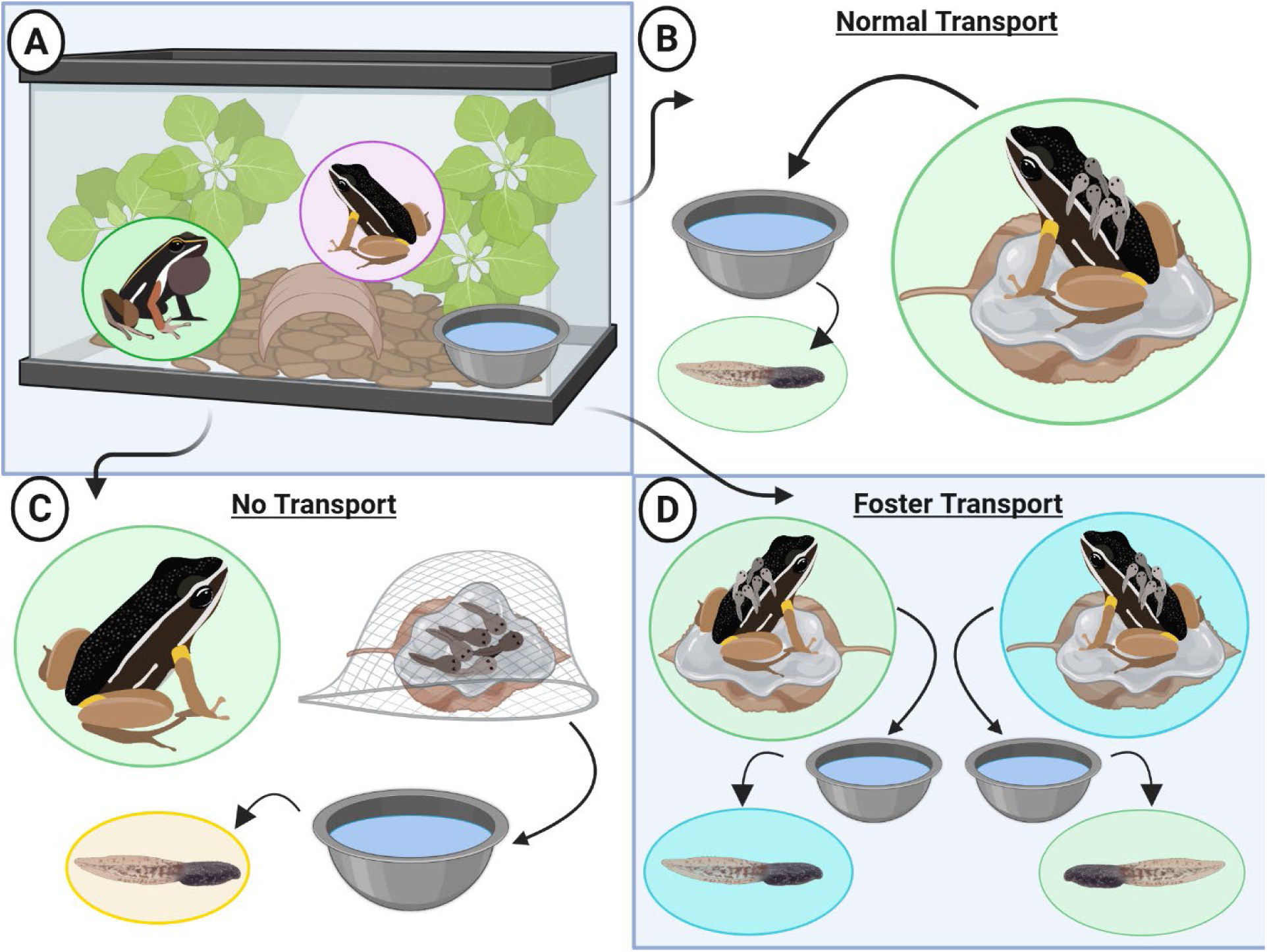
Experimental design. A) Initial setup composition of all trials with adult male and female frogs cohabiting in a tank with substrate (leaves and pebbles), a water bowl and a shelter. The experimental design comprises three treatments, where B) in the ‘normal transport’ treatment, tadpole transport is performed by the biological fathers of the clutches, C) in the ‘no transport’ treatment, tadpoles were prior contact with the father was prevented by placing a net cage over the clutch and tadpoles were manually transferred to the water bowl by experimenters; and D) in the ‘foster transport’ treatment, clutches were exchanged between tanks and tadpoles were transported by foster fathers. Created in BioRender. Angiolani Larrea, F. (2025) https://BioRender.com/5ys3rfw

Our experimental setup allowed us to disentangle microbial acquisition from parents and the environment to offspring over time. We found a clear microbial contribution by the transporting male, regardless of whether the transporting male was the biological or foster father, and that microbial communities were maintained beyond metamorphosis, despite significant differences in relative abundances. The microbial contribution by parents included several bacterial genera that were previously shown to be effective against the detrimental amphibian pathogen *B. dendrobatidis* (*Bd*), highlighting the diverse functional benefits of close parent-offspring interactions.

## Materials and Methods

### Study species and lab population

The Neotropical poison frog *Allobates femoralis* has a pan-Amazonian distribution with isolated local populations. During the reproductive season, males call from elevated structures on the forest floor to announce their territory to male competitors and to attract females [39; 40]. Pair formation, courtship, and mating take place in the male’s territory [41–44], where externally fertilized clutches are deposited in the leaf litter. Tadpole transport to water bodies takes place after 15–20 days of larval development and is mainly performed by males after heavy rains [45]. In the wild*, A. femoralis* uses diverse water bodies ranging from floodplains to medium sized temporal pools for tadpole deposition [40; 46; 47].

We conducted our study in 2022 under controlled laboratory conditions in the animal care facilities at the Ethological station from the University of Bern, Switzerland. Housing conditions are described in the Supplementary Materials.

### Cross-foster experiment and microbiome sampling

We conducted a total of 52 trials, of which 22 could not be considered in the final analysis because of reproduction issues (e.g. the clutch did not hatch or the tadpoles died early in the experiment). Thirty trials were successful; including 23 adult males and 20 adult females in different combinations as breeding pairs. A cleaned and autoclaved water bowl was placed inside the tanks and filled with demineralized water at the start of each trial. Once a breeding pair laid a clutch, we removed the female from the tank and randomly assigned the clutch to one out of three treatments (Fig. 1): In the “normal transport” treatment (N = 9), the male had full access to its own clutch and was allowed to transport the tadpoles to the provided water bodies inside its tank. This treatment was used as a positive control. In the “no transport” treatment (N = 9), a net cage was placed over the clutch to prevent any further physical contact between the male and its clutch. After hatching, the tadpoles were manually transferred to the provided water bowl by placing the leaf where the clutch was deposited. This treatment was used as a negative control. In the “foster transport” treatment (N = 12), we moved clutches together with the leaf they were laid on, to exchange a recently laid clutch with a stage-matched clutch of another breeding pair. New gloves were always used when transferring the clutches or tadpoles to water or between tanks, to avoid cross contamination with human skin microbiome. Adult males in the “foster transport” treatment correspond to both the biological and the foster father. Like the “normal transport” treatment, the foster father had full access to the cross-fostered clutch.

All clutches were closely monitored until hatching. After tadpoles had been deposited in the water bowls (by frogs, or by manual transfer in the “no transport” condition), we poured the water and tadpoles into separate bigger aquaria (20 × 20 × 40 cm) to rear all tadpoles clutch-wise and separate from the tank environment. After metamorphosis (approx. 2 months after transfer), juvenile frogs were moved clutch-wise to rearing enclosures equipped with autoclaved pebbles until final microbiome swabbing (for approx. 2 weeks after metamorphosis).

For microbial sampling, all adults were caught at the onset of each trial with transparent plastic bags, and a new bag and new gloves were used for each frog to avoid cross-contamination. We collected two swabs per individual, one from the ventral abdominal skin and one from the dorsal skin. To obtain information on the environmental microbiome, we swabbed the substrate (leaves and pebbles) inside each tank and dipped swabs in water bowls and lightly scratched the bottom (i.e. tank samples). Adult skin microbiome was sampled a single time to minimize stress of handling for the individuals, even if the same individual was used in different trials (e.g. as biological father in one and foster father in another trial). Microbial samples of clutches were collected after the first (w1) and second (w2) weeks after oviposition by carefully swabbing the surface of the clutch and slightly dipping the swab tip into the egg jelly envelope. Microbial samples of tadpoles were collected by gently swabbing over the main body and tail of tadpoles (up to 5 tadpoles per clutch). Sample collection took place once the tadpoles were big enough to withstand swabbing (at least 1 cm long from mouth to tail tip, corresponding to about 1 month old tadpoles). Finally, we swabbed the dorsum of metamorphs per clutch, once they reached at least 1 cm of snout-vent-length (about 1 week after metamorphosis). All samples were taken using sterile swabs (COPAN, Medical Wire MW100 Swab) and immediately stored at -20°C until DNA extraction.

### DNA extraction, sequencing and bioinformatics

DNA was extracted from swabs using the Macherey-Nagel^TM^ NucleoSpin Soil kit^TM^ (Valencia, CA, USA) according to the manufacturer’s protocol, with lysis Solution 2 (SL2).

We prepared the libraries targeting the 16S rRNA gene at the Genetic Diversity Centre (GDC) from the ETH Zurich through a two-step PCR protocol (see Supplementary Materials for more details). For the first step PCR, we used primers with overhanging adaptors (ITSf: TCGTCGGCAGCGTC-AGATGTGTATAAGAGACAG-[N]3-AG-CCTACGGGNGGCWGCAG; ITSr: GTCTCGTGGGCTCGG-AGATGTGTATAAGAGACAG-[N]3-CC-GACTACHVGGGTATCTAATCC designed at the GDC and adapted from [48]). Demultiplexing, filtering and clustering service was provided by the GDC with the UNOISE pipeline [49] and the SILVA SSU v138 reference was used for taxonomic affiliations.

Data analyses were conducted in R 4.4.0 [50]), with RStudio 2024.04.2 [51]. We used zero-radius Operational Taxonomic Units (zOTUs) for all of our analyses. To determine differential compositions between dorsal and ventral samples from the same frogs, we performed a PERMANOVA with the *adonis_pq* function of the “vegan” package (version 2.6-6.1; [52]) using Bray-Curtis distances and 999 permutations. We did not find compositional differences between ventral and dorsal samples of the same adult frogs (Males: R² = 0.016; p = 0.977; Females: R² = 0.018; p = 0.369). In rare cases (N = 2) we replaced missing dorsal samples with their ventral counterparts.

We obtained valid microbiome information of 22 adult males, 20 adult females, 22 clutches in the first week of development, 25 clutches in the second week of development, 78 tadpoles, 48 metamorphs, 17 substrate (leaves and pebbles), and 21 water (bowl) samples. We use the word “males” to refer to samples from all males used in the entire study; “fathers” is used to refer to either “foster fathers” or “biological fathers” in the different trials.

### Statistical analyses

We quantified differences in microbial composition (beta-diversity) between sample types, calculating Bray-Curtis distances between zOTUs from our samples (“vegan” package) and visualized it with a Non-metric MultiDimensional Scaling (NMDS; Fig. 2A) ordination plot. We did an Analysis of Similarity (ANOSIM) on the calculated Bray-Curtis distances to test for overall differences between sample types followed by pairwise comparisons.

**Figure 2.**
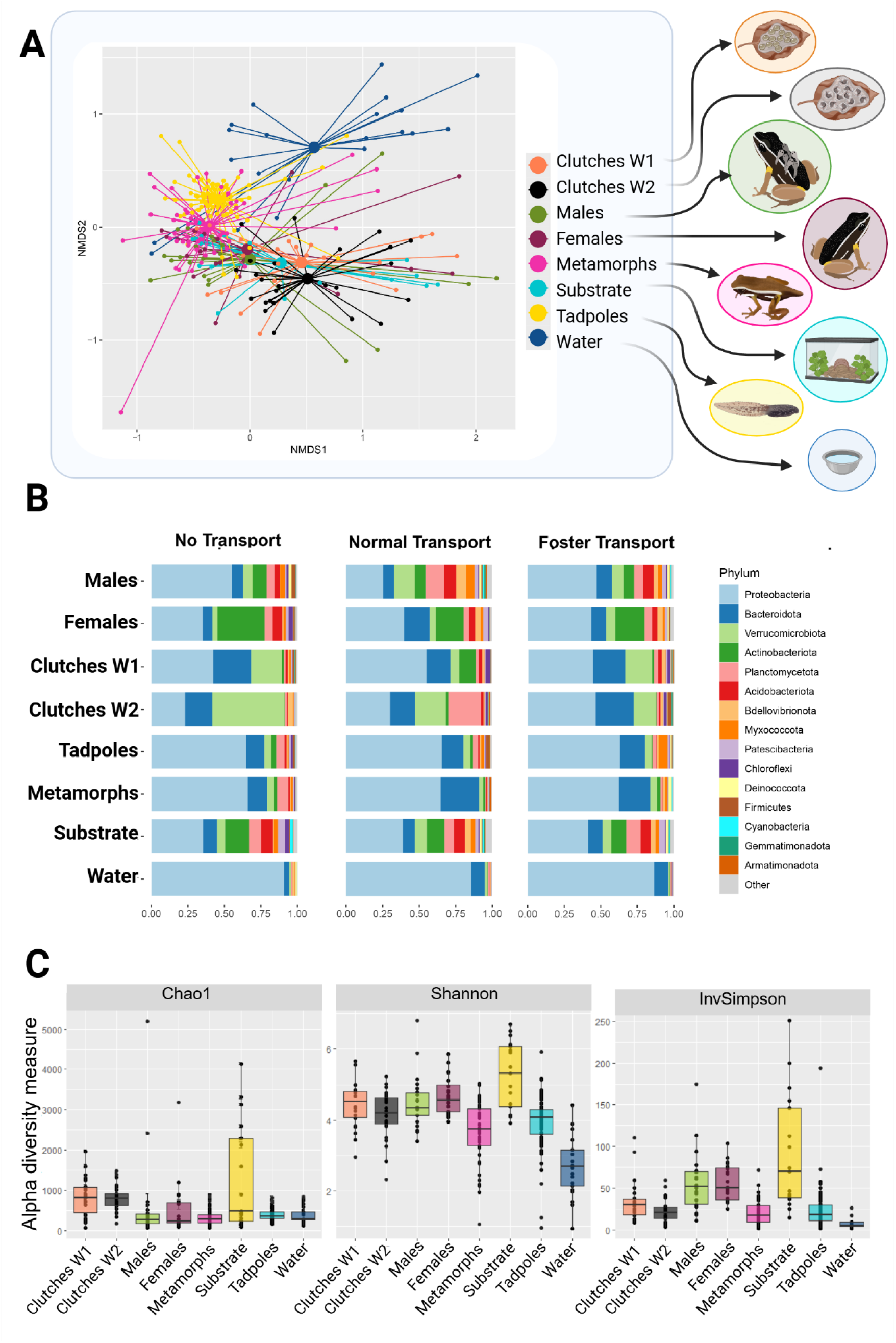
A) Non-metric multidimensional scaling (NMDS) ordination plot of the microbial community found in our experiment clustered by sample types (Bray-Curtis dissimilarity, k = 2, stress = 0.1), B) abundance of microbiome composition by sample types grouped by Phylum, C) calculated alpha diversity indexes (Chao1, Shannon and Inverse Simpson) of the different sample types. Created in BioRender. Angiolani Larrea, F. (2025) https://BioRender.com/cb3pods

To test for differences in microbial alpha-diversity between sample types across treatments, we computed Shannon, Chao1, and Inverse Simpson indexes and tested for differences with either Anova or Kruskal-Wallis tests followed by post-hoc tests when necessary. To evaluate the number of zOTUs that were shared between sample types, we created intersection plots separately by treatment (see Supplementary Materials).

To trace microbial acquisition from the various sources to offspring, we used the Bayesian source-sink tracking tool SourceTracker2 as it considers the entire microbial community profile in complex mixtures [53]. Our different sources were significantly distinct (ANOSIM R = 0.54; *p* = 0.001) further supporting the use of SourceTracker2. To test for differences in the percentage of contribution of each source between experimental conditions, we used a generalized linear mixed effects model with proportion of contribution from sources as response variable, the interaction between treatment, sink and source as fixed effects, and trial identity as random effect. To determine the exact acquisition through tadpole transport, we calculated the zOTUs shared between fathers and tadpoles per experimental condition (normal transport, no transport, foster transport). We also used SourceTracker2 to closely follow the changes in skin microbial composition throughout development. We tested the differences in the proportion of contribution with a generalized linear mixed effects model. We used proportion of contribution from sources as response variable, treatment and sink as fixed effects, and trial identity as a random effect.

To identify which putative *Bd* inhibitory taxa were acquired by the offspring from adults during tadpole transport, we first inspected our full microbiome dataset to a published list of taxa known to inhibit the fungus *Bd* [7]. We then compared the zOTUs within tadpole and metamorph samples for each treatment to this list. To determine if the acquisition of putative Bd inhibitory taxa was a non-random process, we used generalized linear models to compare the overall count of shared zOTUs by treatment for tadpoles and metamorphs separately. We used the number of putative Bd inhibitor taxa shared between fathers and offspring per set, as the response variable, and treatment as the independent variable. We diagnosed our models and tested for significance of independent variables with an Analysis of Deviance (Type II Wald chi square test), followed by paired comparisons. More details on the methods are provided in the Supplementary Material.

## Results

We found significant differences in microbial composition across sample types (ANOSIM R = 0.54; *p* = 0.001; Fig. 2A, 2B). The pairwise comparisons confirmed that most sample types differed significantly from each other (*p* <0.05, see Supplementary Materials for the detailed results). However, we did not find significant differences in microbial composition between male and female adult samples (ANOSIM R = -0.03; *p-adj* = 1), nor between adult samples and the substrate (males vs. substrate: ANOSIM R = 0.05; *p-adj* = 1; female vs. substrate: ANOSIM R = 0.02; *p-adj* = 1). Moreover, the microbial composition of clutches from week 1 and males (ANOSIM R = 0.28; *p-adj* = 0.06) and of females and metamorphs (ANOSIM R = 0.26; *p-adj* = 0.06) did not differ significantly from each other. A detailed description of the microbial composition of different sample types is given in the Supplementary Materials.

When looking at overall alpha-diversity, we found significant differences between sample types (Kruskal-Wallis, Chao1 *X^2^* = 57.68, df = 7, *p* < 0.01; Shannon, X*^2^* = 85.34, df = 7, *p* < 0.01; Inverse Simpson X*^2^* = 102.14, df = 7, *p* < 0.01; Fig. 2C). Therefore, we explored pairwise comparisons between sample types in which substrate and adult samples were not significantly diverse from each other. Adult male and female samples also did not differ significantly from each other. Likewise, tadpole and metamorph microbial diversity did not differ. Additionally, we found no significant differences of the effect of treatment on the alpha-diversity of offspring samples (see Supplementary Materials for more details).

### Sources of microbial acquisition

We found a higher number of shared zOTUs between the adult and the substrate samples (normal transport: 995/97 (male/female) shared zOTUs; no-transport: 63/105 shared zOTUs; foster transport: 188/102 shared zOTUs, see also Supplementary Materials) than the rest of all possible combinations. Maternal samples shared 18 zOTUs with tadpole samples in the normal transport treatment, 24 in the no-transport treatment, and 12 for the foster treatment. Adult male samples had no detectable shared zOTUs with the tadpoles or metamorphs in the no-transport treatment. In turn, there were clearly detectable shared zOTUs in the normal transport (tadpoles: 37 shared zOTUs; metamorphs: 15 shared zOTUs) and foster transport (tadpoles: 21 shared zOTUs; metamorphs: 11 shared zOTUs) treatments. We found the highest overlap of taxa between tadpoles and tank samples in the no-transport treatment (substrate: 160 shared zOTUs; water: 79 shared zOTUs), while the normal transport (substrate: 17 shared zOTUs; water: 35 shared zOTUs) and the foster transport treatment (substrate: 31 shared zOTUs; water: 10 shared zOTUs) had considerably lower amounts of shared taxa with the tank samples.

The SourceTracker analysis identified that substrate, water, and both parents were all important sources for the acquisition of skin microbiota in offspring (Fig. 3, Table S1). However, we found that neither treatment, source or the interaction between treatment, source, and sink (i.e. offspring) revealed significant differences in relative contribution of microbial taxa across developmental stages (Fig. 4A, Table S2). However, we found significant differences in relative contribution of the various sources between sinks (Anova.glmmTMB *X^2^* = 13.23, df = 3, *p* = 0.004). We found that a large part of the microbiome was changing between developmental stages, from clutches to tadpoles and metamorphs regardless of treatment (Fig. 4B, treatment: Anova.glmmTMB *X^2^* = 1.58, df = 2, *p* = 0.455; developmental stage: Anova.glmmTMB *X^2^* = 80.45, df = 2, *p* < 0.001). We found that hatching was the stage with significantly less microbial contribution from source to sink (embryonic development - hatching: β = 1.66, SE = 0.23, z = 7.40, *p* < 0.001; hatching - metamorphosis: β = -1.40, SE = 0.18, z = -7.78, *p* < 0.001; embryonic development - metamorphosis: β = 0.25, SE = 0.22, z = 1.16, *p* = 0.481). The change in microbial composition during embryonic development was most variable across treatments but relatively high compared to all other stages (contribution from clutches w1 to clutches w2: normal transport: 29.24% ± 22.86; no transport: 56.6% ± 28.98; foster transport: 33.54% ± 26.9). After hatching, the microbiome composition shifted considerably, as the microbial contribution of clutch w2 to the tadpole stage was very low (normal transport: 4.37 % ± 3.5; no transport: 7.95 % ± 12.27; foster transport: 8.57 % ± 9.39, Fig. 3). In contrast, the maintenance of the microbiome from tadpoles to metamorphs was high (normal transport: 32.13 % ± 20.87; no transport: 34.95 % ± 16.01; foster transport: 32.69 % ± 17.93, Table S1). We identified significant deviations from model assumptions (overdispersion: *p* = 0.016; uniformity: *p* = 0.014), however, this can be explained by the large sample size of our dataset and the sensitivity of the DHARMa test [54]. Visual inspection of the QQ plot residuals suggest that the distribution of residuals is very close to the expected optimal values (see Supplementary Materials).

**Figure 3.**
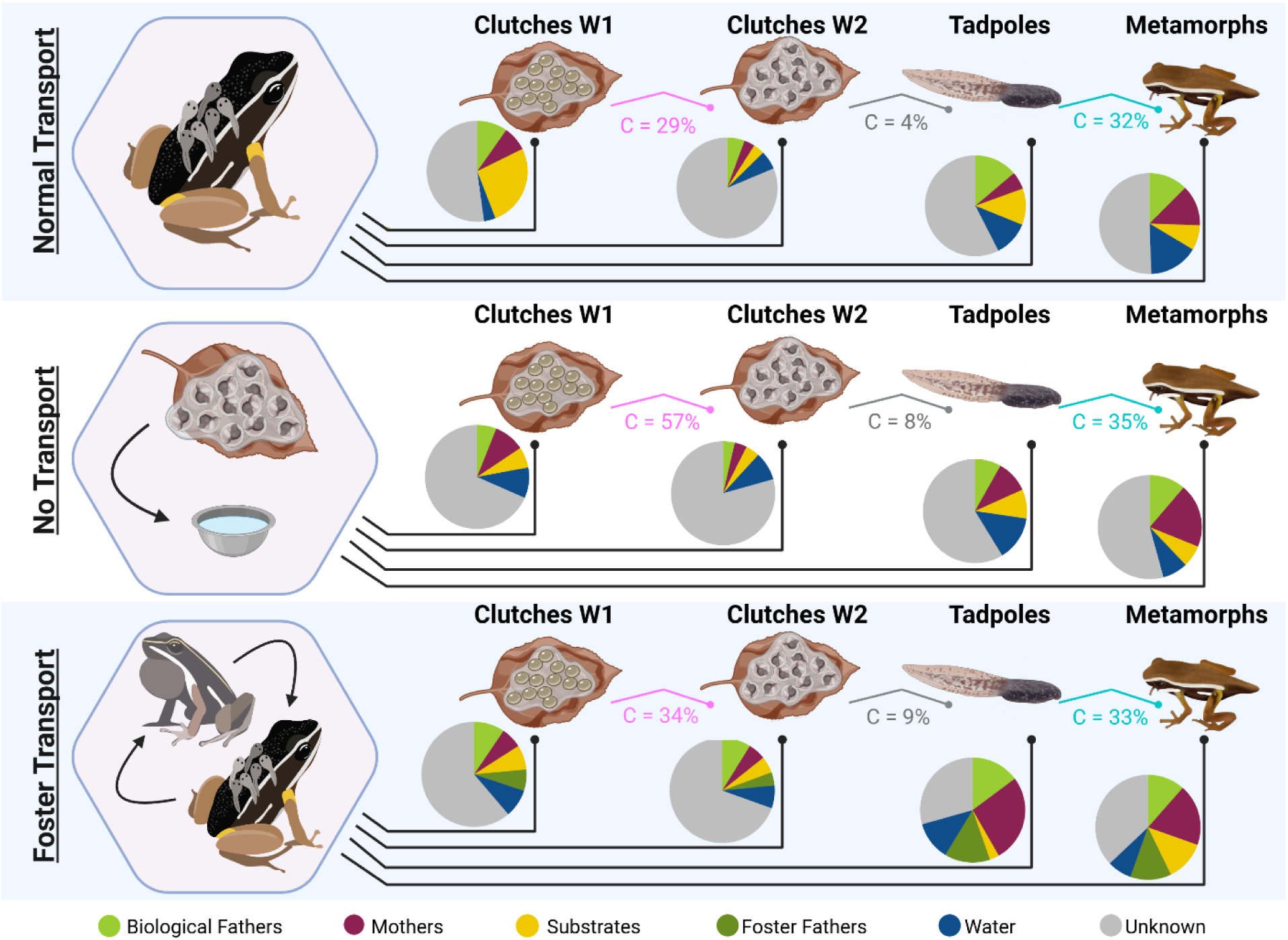
Relative contributions of all tested sources to the different developmental stages of the offspring determined by the SourceTracker2 analysis. Pie charts show the microbial contribution from each source to the respective sinks. C = contribution (%) from source to sink. Created in BioRender. Angiolani Larrea, F. (2025) https://BioRender.com/5qgkbk9

**Figure 4.**
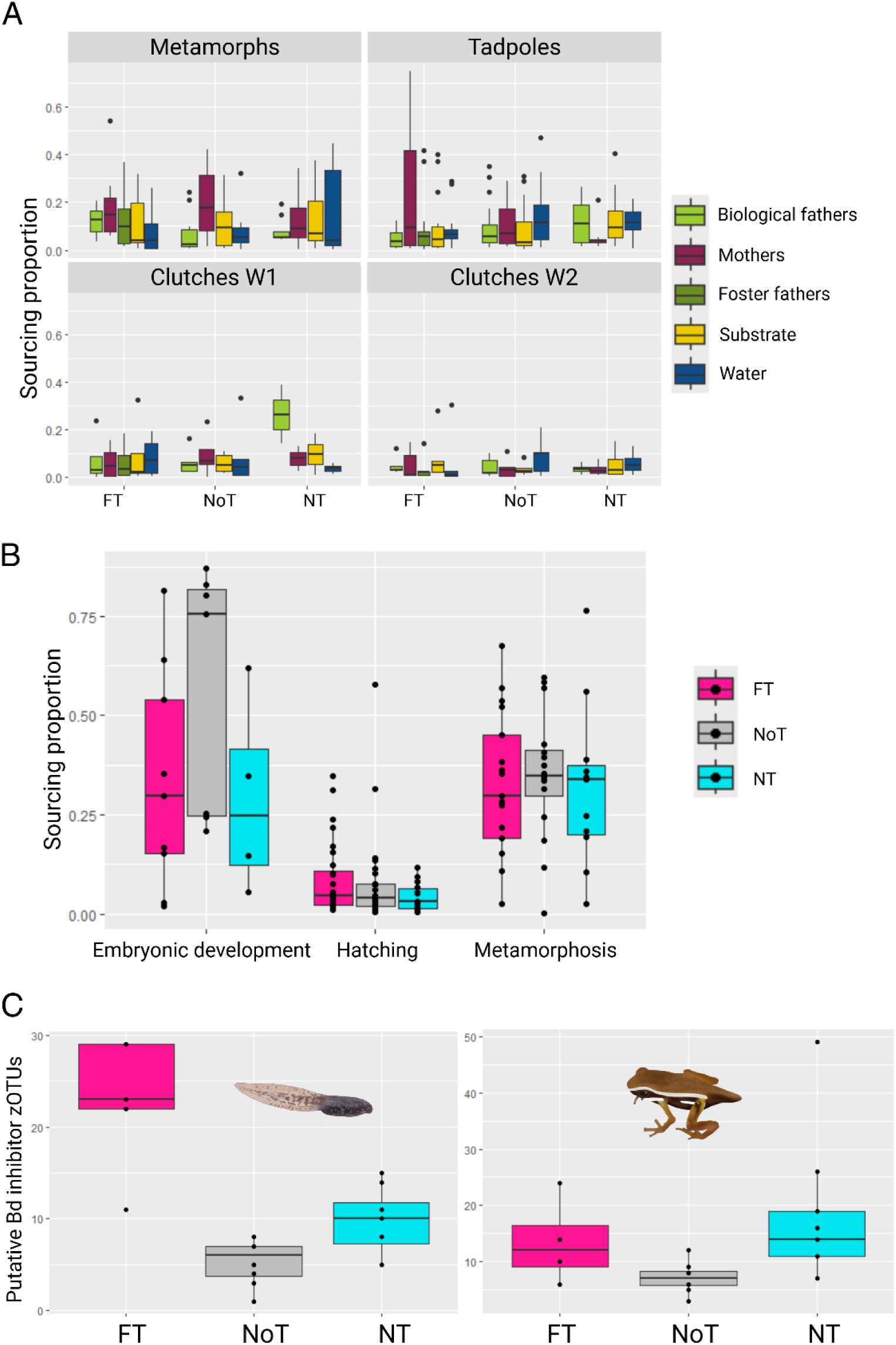
A) Sourcing proportions of acquired zOTUs per tested source by offspring sample and treatment. B) Sourcing proportions across developmental stages per treatment. C) Number of zOTUs of putative Bd inhibitory taxa shared between foster father (foster transport), non-transporting biological father (no transport) and transporting biological father (normal transport) with tadpoles (left panel) and metamorphs (right panel). FT = Foster transport, NoT= No transport, NT = Normal transport. Created in BioRender. Angiolani Larrea, F. (2025) https://BioRender.com/6euq3ym

### Inhibitory taxa acquired

Using a presence–absence approach, we identified zOTUs shared exclusively between adult males and their offspring. In the normal transport treatment, biological fathers shared 165 zOTUs with their tadpoles (including 32 putative Bd-inhibitory taxa) and 289 zOTUs with their metamorphs (69 inhibitors). In the no-transport treatment, non-transporting fathers shared 72 zOTUs with tadpoles (16 inhibitors) and 91 zOTUs with metamorphs (26 inhibitors). In the foster transport treatment, foster fathers shared 107 zOTUs with tadpoles (27 inhibitors) and 143 zOTUs with metamorphs (33 inhibitors). Within the foster transport treatment, at least 16 putative Bd-inhibitory taxa persisted from tadpoles to metamorphs, spanning the genera *Pseudomonas, Comamonas, Chryseobacterium, Acidovorax, Aeromonas, Staphylococcus, Shewanella, Sphingomonas, Brevundimonas,* and *Acinetobacter,* as well as *Sphingobacterium spiritivorum* and *Herbaspirillum* sp. No dominant putative Bd-inhibitory zOTUs were detected in water samples.

Treatment had a significant effect on the total number of putative *Bd* inhibitor zOTUs shared between adult males and offspring (tadpoles: Anova *X^2^* = 79.19, df = 2, *p* < 0.001; metamorphs: Anova.glmmTMB *X^2^* = 13.31, df = 2, *p* = 0.001; Fig. 4C). In tadpoles, we found that all three treatments differed significantly from each other, with the no transport treatment having the lowest number of shared zOTUs compared to the other treatments (foster transport - no transport: β = 1.47, SE = 0.18, z = 8.14, *p* < 0.001; foster transport - normal transport: β = 0.85, SE = 0.15, z = 5.78, *p* < 0.001; normal transport - no transport: β = -0.62, SE = 0.19, z = -3.23, *p* = 0.004; Fig. 4C). In metamorphs, the results show a similar pattern, as also the no transport condition showed the lowest number of shared zOTUs. In the no transport treatment, the shared zOTUs of putative *Bd* inhibitors with adults was significantly lower than in the normal transport treatment (normal transport - no transport: β = -0.94, SE = 0.26, z = -3.64, *p* < 0.001), but none of the other comparisons revealed statistically significant differences (foster transport - no transport: β = 0.64, SE = 0.32, z = 1.98, *p* < 0.117; foster transport - normal transport: β = -0.30, SE = 0.30, z = -1, *p* = 0.578; Fig. 4C).

## Discussion

We examined the role of social interactions during parental care for the acquisition of key skin microbiota in the poison frog *Allobates femoralis*. Using a cross-foster experimental design, we provide causational evidence that tadpole transport is an important source for microbial seeding in offspring. Some of these acquired microbiota are putatively inhibitory to the detrimental chytrid fungus *B. dendrobatitis.* Further, the microbiome composition is largely retained beyond metamorphosis, including in particular putative protective bacteria. These findings provide novel insights into the multifaceted roles of parental care for equipping the next generation with essential resources, such as diverse and targeted skin microbial communities.

In our study, contact with the clutch during oviposition and fertilization by both adults appeared to provide microbial seeding strong enough to preserve parental microbial signatures throughout development from eggs to metamorphs. Despite removing the mother from the tanks when starting a trial, the mother’s microbiome signature was still detected in offspring, from clutches to metamorphs, across all treatments. Our finding illustrates that some initial microbial seeding by mothers is taking place already before or during oviposition. In poison frogs, the vitelline membrane has been described as a barrier to microorganismal exchange between embryos and the jelly [31], rendering the embryos virtually sterile while inside the eggs. However, upon hatching, embryos immediately come into contact with the jelly matrix until the caring parent (male) collects the hatchlings. This interaction exposes the hatchlings to the jelly-associated microbiota, initiating colonization of the tadpole skin [31]. These newly inoculated microbes have the potential to then become stable colonizers on the offspring skin, as supported by our study.

Similar to maternal contributions, males seem to contribute to the microbial seeding already during fertilization. The biological father’s microbial signature was found in offspring in the foster transport treatment, and in the no transport treatment, and these signatures remained traceable beyond metamorphosis. However, a considerable amount of microbiota is also acquired during tadpole transport. In the foster transport treatment, the microbiome of the foster father did not fully replace that of the biological father, nor did the initial seeding by the biological father prevent additional microbial acquisition from the foster father (Fig. 3). Hence, our results demonstrate that vertically transmitted microbiota play both seeding and reinforcing roles in microbial community establishment (see also [55] for an example in gut microbiota). Moreover, the increase in the percentage of contributions of microbial communities by adults after metamorphosis suggests a priority effect of these initially acquired microbiota [56].

An important source of microbial acquisition in animals is the environment (e.g., [32; 57; 58]. The lack of statistical differences in microbial composition between adult frog and substrate samples are probably due to captivity [59]. However, growing evidence suggests that social interactions, including parental care, can also impact the microbiome. Previous studies have demonstrated that close social interactions can even reduce the contribution of the environment through e.g. engraftment processes (amphibians: [16; 28; 31]; fish: [60]). Our results are in line with these findings, as the number of shared taxa between the environment and offspring was considerably lower in the two conditions involving transport by male frogs (natural transport and foster transport) compared to the ‘no transport’ condition. We assume that the males transferred beneficial microbes to the tadpoles that subsequently engrafted on the skin.

We identified important shifts in microbial composition across offspring development. Hatching (i.e. the first contact of tadpoles with the external environment) was associated with major shifts in skin microbiome composition. The contribution of previously seeded microorganisms from known sources acquired during embryonic development dropped to 4–9% post-hatching. Hatching is a critical step for microbiota acquisition in egg-laying animals, as larvae transition from relatively stable embryonic conditions to novel environments. Novel contact with environmental microbial sources has been suggested as a strong filter for microbial composition of the host, potentially enhancing the establishment of parental colonizers [55]. In turn, we found that the microbial contribution from tadpoles to metamorphs was significantly higher (32-35%) than in the pre-hatching stages. These shifts, together with the non-significant differences in alpha diversity between tadpole and metamorph samples, suggest recurring recruitment of microbial communities that established and grew during development. The observed differences between tadpole and metamorph samples, suggest that a reshaping process of the microbial community takes place during metamorphosis. Changes in microbial community composition over ontogeny are widespread among animals (e.g., humans: [13; 61]; non-human mammals: [62]; birds: [63]; reptiles: [64]; amphibians [32; 65]; fish: [14; 60]; insects: [19; 66]; other invertebrates: [67]. Major shifts in environmental conditions offer numerous opportunities for colonization by new bacteria. Colonization may however, be influenced by priority effects acting either by competition (as evidenced in our experiment after hatching) or facilitation (as evidenced in our experiment after metamorphosis) with previously acquired microorganisms (reviewed by [68]; e.g., amphibians: [28; 31; 59; 69]; fish: [70]; insects: [55; 71; 72]). Metamorphosis in amphibians represents such a major morphological and physiological transition from aquatic to terrestrial environments that translate to major changes in microbiome richness and composition [32]. In our study, we identified a considerable reorganization of microbial communities across egg to metamorph development. However, the initial seeding by parents was traceable across the development, and tadpole transport reinforced the maturation of the offspring skin microbiome.

Social interactions have been shown to facilitate the colonization of specialized taxa [71]. We observed a similar effect in our study: a substantial portion of the acquired taxa includes bacteria previously identified as key components in the immune response against *Bd* — a globally emerged pathogenic fungus [7; 20]. We acknowledge that in our analysis we did not identify the specific strains that correspond to known Bd inhibitory taxa and we remain at the genus level. These taxa may still include bacterial taxa that are not specifically inhibitors or may even facilitate infection by Bd [73]. The acquired potentially protective bacteria persisted even past metamorphosis, though the proportions shifted, indicating succession processes founded in social interactions within the context of parental care [1; 74]. Our findings underline the resilience of these potentially beneficially microbial communities during extreme developmental transitions in offspring, such as tadpoles hatching in terrestrial environments and undergoing metamorphosis [75].

## Conclusions

Our study provides clear experimental evidence for vertical transmission of skin microbiota during parental care in a poison frog, and the retention of these microbial communities beyond metamorphosis, despite significant differences in relative abundances. In poison frogs, tadpole transport is an obligate behaviour, critical for the survival of tadpoles hatched in terrestrial environments; by actively relocating larvae to water, parents prevent desiccation and ensure offspring survival (*28, 35*). Our findings suggest that tadpole transport is more than just a behavioural adaptation to ensure the survival of terrestrial eggs that develop into aquatic larvae. Our findings suggest that beneficial microbiota can be acquired through close parent-offspring interactions, are sustained over metamorphosis, and have the potential to increase the offspring’s ability to withstand detrimental diseases. Understanding the role of parental care in enhancing social immunity could provide key insights into the mechanisms that have led to the evolution of parental care in animals.

## Acknowledgements

We want to thank David Johnson and the staff from the GDC, Silvia Kobler, Aria Minder, Jean-Claude Walser and Niklaus Zemp for their assistance and guidance during sample preparation and processing. We also want to thank Christoph Netz for his help on statistical analyses. Finally, we want to thank Evi Zwygart, Anyelet Valencia-Aguilar, Marina Garrido-Priego and Lauriane Bégué for their help with animal care and assistance during microbial sampling.

## Data availability statement

Sequences have been deposited in the “Inventaire des Données de la Recherche et Environnement et Sociétés” (InDoRES) repository, (link will become available once the paper is accepted)

Code and further raw data used are available here: (will be converted into a stable link after acceptance)

## Author contributions

FNAL: Conceptualization, Data Curation, Formal Analysis, Investigation, Methodology, Visualization, Project administration, Writing – Original Draft Preparation;

RB: Formal Analysis, Visualization, Data Curation, Methodology, Writing – Review & Editing;

AL: Conceptualization, Methodology, Visualization, Funding Acquisition, Validation, Writing – Review & Editing;

DS: Conceptualization, Resources, Methodology, Validation, Supervision, Writing – Review & Editing;

ER: Conceptualization, Funding Acquisition, Methodology, Supervision, Resources, Writing – Original Draft Preparation, Project administration, Writing – Review & Editing

## Ethics approval statement

All testing was approved by the Swiss Federal Food Safety and Veterinary Office (National No. 33232, Cantonal No. BE144/2020). Captive conditions were approved by the Swiss Federal Food Safety and Veterinary Office (Laboratory animal husbandry license: No. BE4/11). We followed the guidelines laid out by the ASAB for the treatment of animals in behavioural research and Teaching [76] and the ARRIVE guidelines [77].

The founding individuals of this population were sampled in and exported from French Guiana in compliance with all legal requirements from the responsible French authorities (DIREN: Arrêté n°82 du 10.08.2012 and Arrêté n°4 du 14.01.2013). Individuals used in the present study are all captive bred individuals from this original stock.

## Further declarations

We used ChatGPT and Deepseek for language editing. We used BioRender for creating the figures.

## Supplementary materials

### Detailed methods section

#### Laboratory frog population – housing conditions

Individuals were kept in breeding pairs in standard (60 × 40 × 40 cm) glass terraria furnished with a coconut shelter, a perch, a plant, a water bowl, and expanded clay pebbles covered with autoclaved oak leaves. The sides were covered with Xaxim (tree fern stems) and cork mats to prevent visual contact between terraria. Light, temperature (daily cycles between 23-28°C), and hydration of tanks (rain cycles twice per day to ensure humidity close to 100%) were automatically controlled to mimic natural conditions in French Guiana. All frogs were fed twice a week with vitamin-dusted fruit flies. Tadpoles received dried *Urtica* leaves ad libitum and were kept in aquaria filled with reverse osmosis water, equipped with a standard aquarium filter (Amazonas Aqua Cleaner with activated carbon and sponge filter) and air supply.

#### DNA extraction PCR conditions

We used the KAPA HiFi DNA Polymerase Kit for the PCR steps. In the first step PCR, conditions were: 3 minutes at 95° C, 20 seconds at 95°C, 15 seconds at 54°C, 15 seconds at 72°C and 5 minutes at 72°C in triplicates for each sample, which were pooled afterwards. Given the low yield of some samples, we pooled amplifications products from up to three PCR runs before sequencing. We used Beckam Coulter Life Sciences Agencourt AMPure XP Beads (Indianapolis, Indiana, USA) to retrieve clean DNA from the PCR product. In the second PCR step, we used the Illumina Nextera XT DNA library preparation Kit (San Diego, CA, USA) following the kit’s protocol specification and 55°C as annealing temperature. Amplicons were pooled, cleaned with beads, and sequenced on an Illumina MiSeq instrument at the GDC.

#### Bioinformatics

Data analyses were conducted in R 4.4.0 ^1^, with RStudio 2024.04.2 ^2^. We first built a phyloseq object, using the package “phyloseq” (version 1.48.0; ^3^) and the *tax_fix* function in “MicroViz” (version 0.12.3; ^4^) to fill empty taxonomic information with the higher known taxonomic rank. We used zero-radius Operational Taxonomic Units (zOTUs) for all of our analyses and zOTUs belonging to non-determined eukaryotic (N = 2), mitochondria family (N = 27), chloroplast class (N = 41) and Archaea taxa (N = 41) were removed.

We also discarded zOTUs with less than 1000 reads. Additionally, we eliminated contamination from our samples with the package “decontam” version 1.24.0 identifying contaminants with a threshold of 0.1 using negative control samples as reference ^5^. We detected 0.12% of contamination corresponding to 27 zOTUs which were removed from the dataset. After quality control, our data set was composed of 253 samples and we had a total of 15 047 092 reads with an average of 59 474 67 (min: 1 080; max: 1 854 976) reads clustered into 8 864 zOTUs.

To determine differential compositions between dorsal and ventral samples from the same frogs, we performed a PERMANOVA with the *adonis_pq* function of the “vegan” package (version 2.6-6.1; ^6^) using Bray-Curtis distances and 999 permutations. We did not find compositional differences between ventral and dorsal samples of the same adult frogs (Males: R² = 0.016; p = 0.977; Females: R² = 0.018; p = 0.369). In rare cases (N = 2) we replaced missing dorsal samples with their ventral counterparts.

We obtained valid microbiome information of 22 adult males, 20 adult females, 22 clutches in the first week of development, 25 clutches in the second week of development, 78 tadpoles, 48 metamorphs, 17 substrate (leaves and pebbles), and 21 water (bowl) samples.

#### Statistical analyses

We quantified differences in microbial composition (beta-diversity) between sample types, calculating Bray-Curtis distances between zOTUs from our samples (“vegan” package) and visualized it with a Non-metric MultiDimensional Scaling (NMDS) ordination plot using the wrapping function *metaMDS* (“vegan” package). We did an Analysis of Similarity (ANOSIM) on the calculated Bray-Curtis distances to test for overall differences between sample types with the function *anosim* (“vegan” package) and 999 permutations. Consecutively, we did pairwise comparisons with the function *anosim.pairwise* (“VeganEx” package, version 0.1.0; ^7^). We applied a Bonferroni correction to account for multiple comparisons.

To test for differences in microbial alpha-diversity between sample types across treatments, we computed Shannon, Chao1, and Inverse Simpson indexes (“phyloseq” function *estimate_richness*) and performed an Anova (function *aov* of basic R) or a Kruskal-Wallis test (function *kruskal.test* from the package “phyloseq”), depending on the distribution of the index values. If the Kruskal-Wallis test revealed significant differences between groups, we performed pairwise comparisons (function *pairwise.wilcox.test* from the package “phyloseq”). We applied a Bonferroni correction to account for multiple comparisons.

To evaluate the number of zOTUs that were shared between sample types, we first created intersection plots separately by treatment (Figure S1) using the functions *upset_set_size* of the “ComplexUpset” package (version 1.3.3; ^8^) and *upset_pq* of the “MiscMetabar” package (version 0.10.1; ^9^). For readability when computing intersections, we set the minimum number of reads per zOTU found per sample to 10, and the minimum set size to 7.

**Figure S1.**
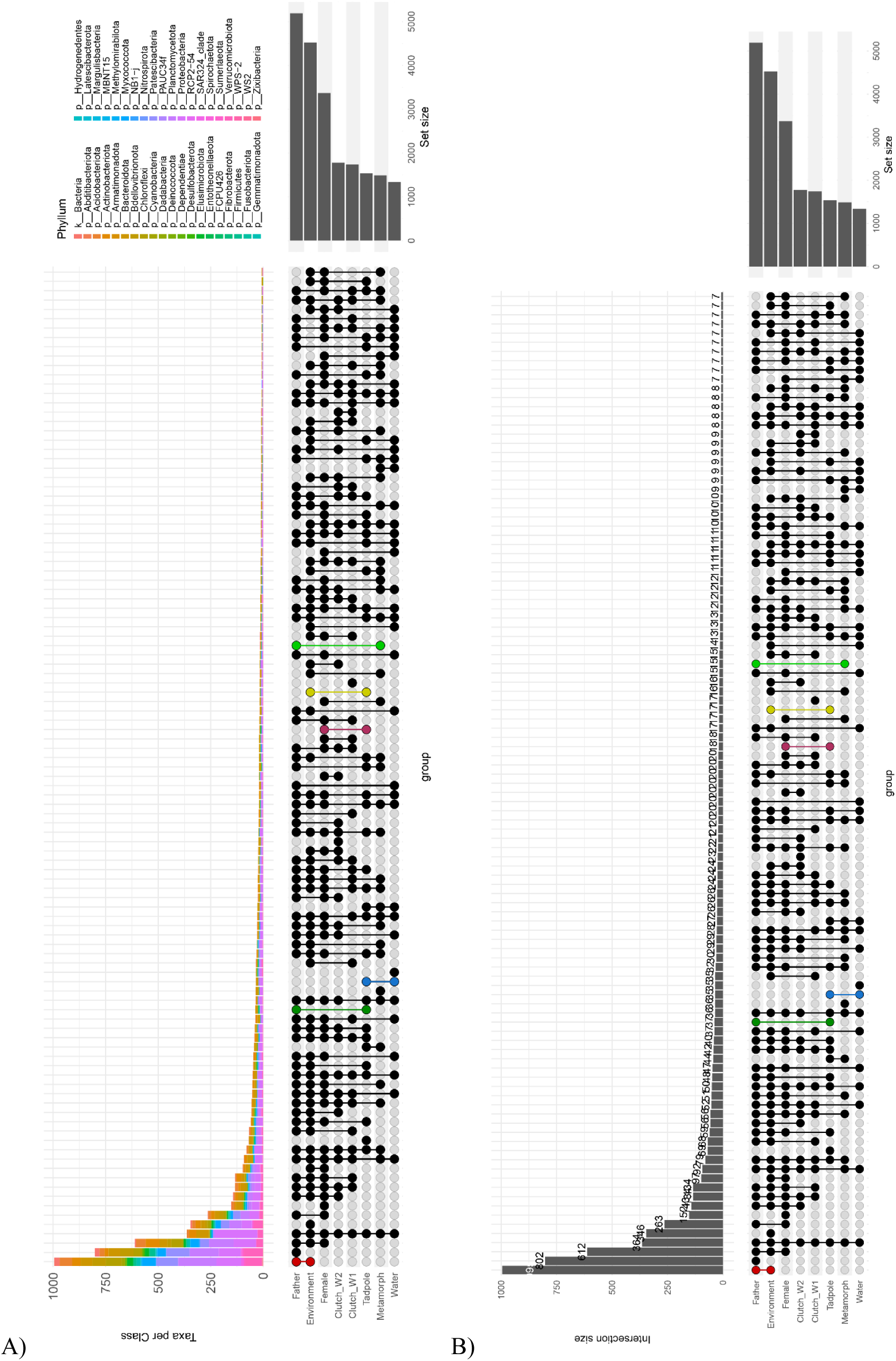
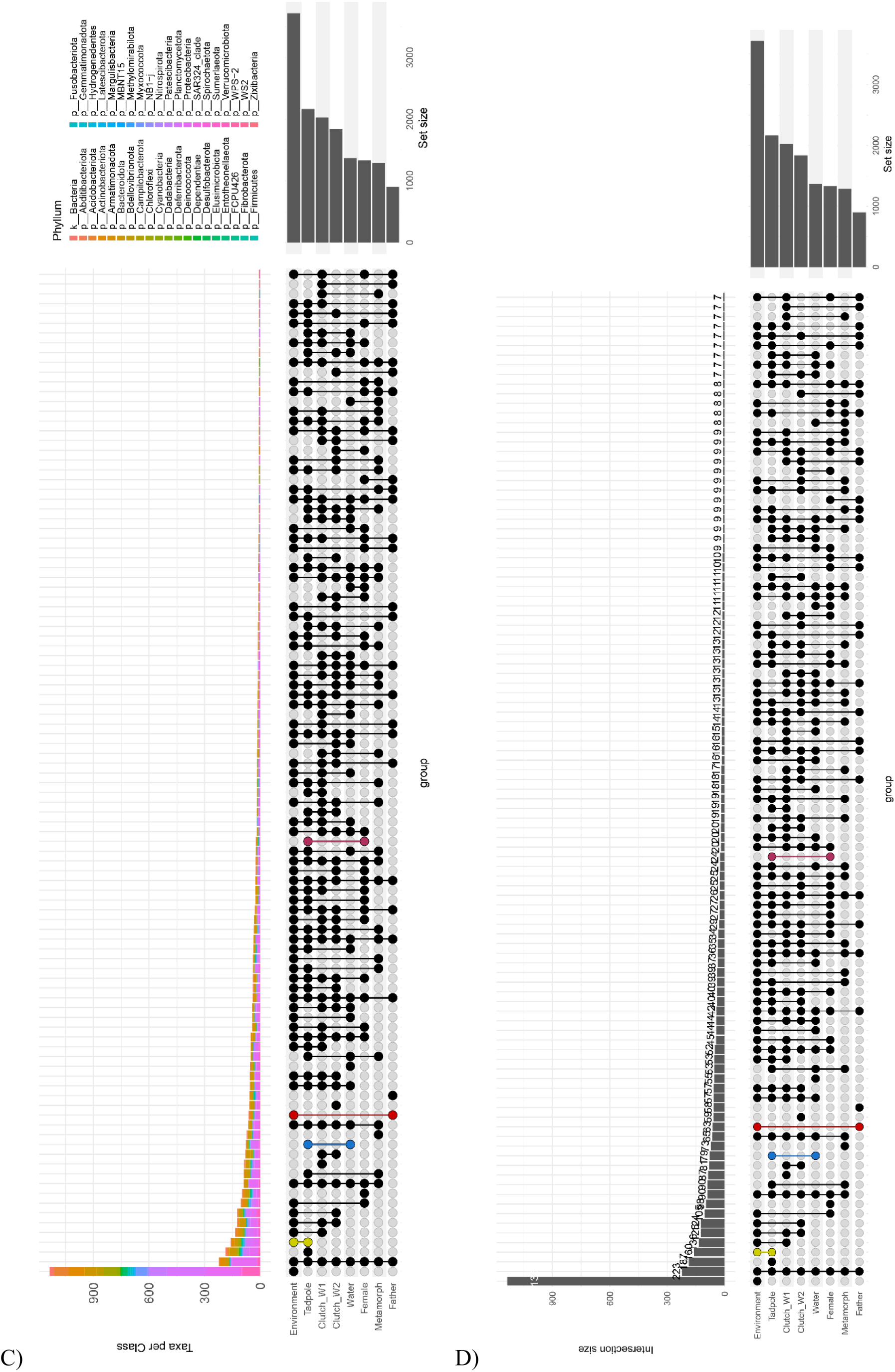
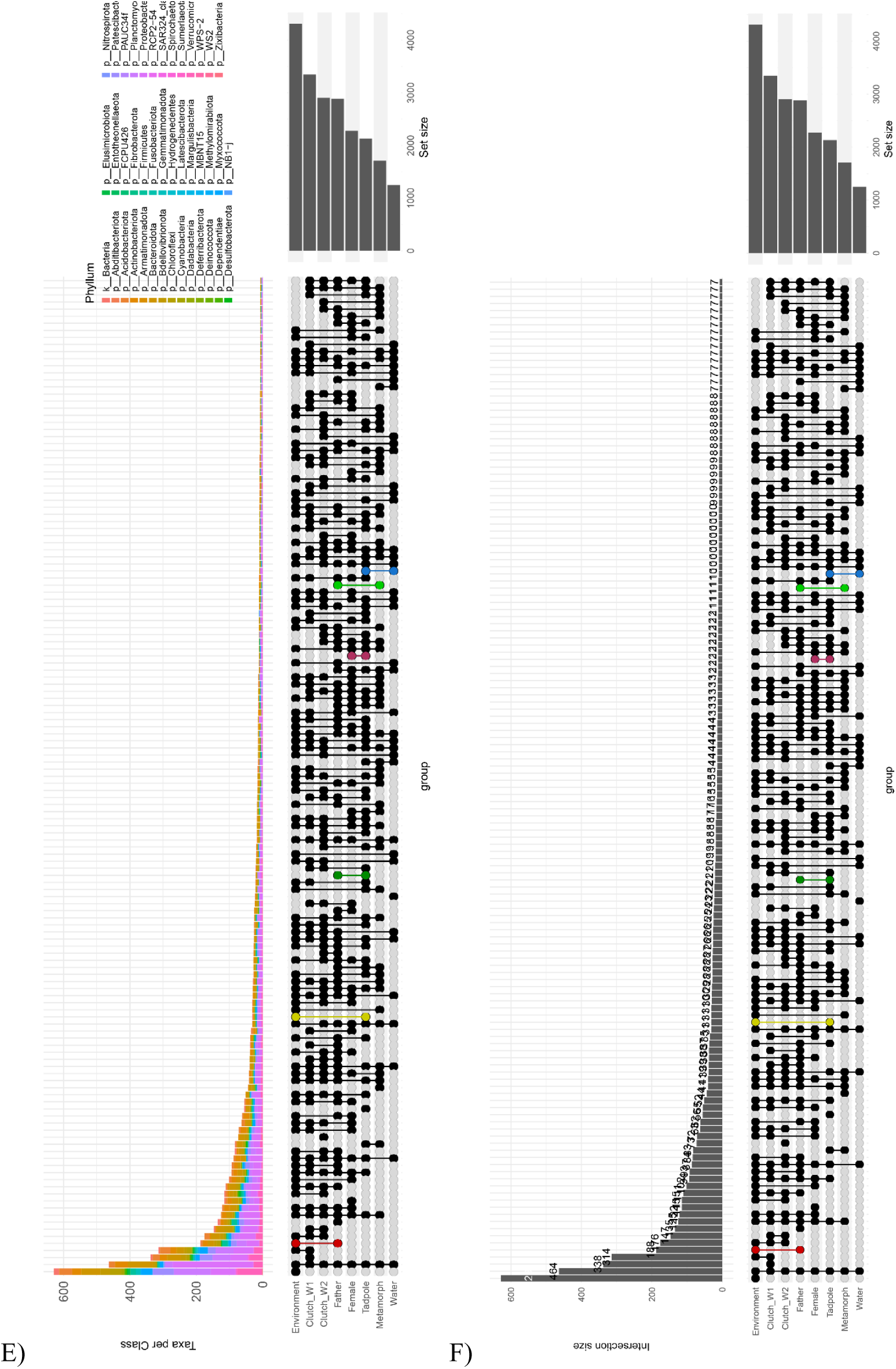
Intersection plots. A) Normal Transport with taxonomic composition at the Phylum level. B) Normal Transport with intersection sizes. C) No Transport with taxonomic composition at the Phylum level. C) No Transport with intersection sizes. E) Foster Transport with taxonomic composition at the Phylum level. F) Foster Transport with intersection sizes.

To trace microbial acquisition from the various sources to offspring, we used the Bayesian source-sink tracking tool SourceTracker ^10^. We performed all SourceTracker analyses with the default parameters (rarefaction = 1 000 reads; burn-in = 100; restart = 10) and ran the analysis three times to account for stochastic effects. We included the corresponding biological father, foster father (in the case of foster transport treatment), mother, substrate, and water as sources, and each different developmental stage of offspring (i.e. clutches week 1 and 2, tadpoles, or metamorphs) as a sink. This analysis was conducted separately for each trial, and in the case of tadpoles and metamorphs for each individual sample. We calculated the mean and standard error for sinks (Table 1). To test for differences in the percentage of contribution of each source between experimental conditions, we used a generalized linear mixed effects model using the *glmmTMB* function (“glmmTMB” package, version 1.1.10 ^11^) and a beta distribution with proportion of contribution from sources as response variable, the interaction between treatment, sink and source as fixed, and trial identity as random effect.

To determine the exact acquisition through tadpole transport, we calculated the zOTUs shared between fathers and tadpoles per experimental condition (normal transport, no transport, foster transport). For this, we used the package “metagMisc” (version 0.5.0; ^12^) and performed the analyses for each trial separately. First, we isolated the zOTUs of the tadpoles in one phyloseq object and all sources (mother, father, substrate, water) in a second one. From the sources, we then extracted the zOTUs that were found only in the fathers with the function *phyloseq_extract_non_shared_otus*. For the foster transport treatment, we extracted the zOTUs that were unique to the foster father (from here on foster father’s unique zOTUs). Then, after merging both phyloseq objects, we identified the zOTUs that were shared between the fathers and the tadpoles with the function *phyloseq_extract_shared_otus* (Supplementary Material S2). To determine if the acquired taxa are sustained over time, we repeated the operation to obtain the shared zOTUs between fathers and metamorphs.

To closely follow the changes in skin microbial composition throughout development, we calculated the proportional contribution from one developmental stage to the next using SourceTracker. We defined three steps: 1) Embryonic development, where clutch week 1 was used as source and clutch week 2 as sink; 2) Hatching, where clutch week 2 was used as source and tadpoles as sink; and lastly 3) Metamorphosis, where tadpoles were used as source and metamorphs as sink (Table 1). We tested the differences in the proportion of contribution by applying a generalized linear mixed effects model using the *glmmTMB* function. We used proportion of contribution from sources as response variable, and treatment and sink as fixed effects. We used trial identity as a random effect and a beta distribution.

To identify which putative *Bd* inhibitory taxa were acquired by the offspring from adults during tadpole transport, we first inspected our full microbiome dataset to a published list of taxa known to inhibit the fungus *Bd* ^13^. We then compared the zOTUs within tadpole and metamorph samples for each treatment to this list (Supplementary Material S2). For this analysis we used only the trials that had microbiome sequence data of all associated samples, adult males, adult female, substrate and water, available (tadpoles: normal transport: N = 8; no transport: N = 8; foster transport: N = 5; metamorphs: normal transport: N = 9; no transport: N = 8; foster transport: N = 4). Additionally, to determine if the acquisition of putative *Bd* inhibitory taxa was a non-random process, we used generalized linear models (GLMs; *glm* function from basic R) to compare the overall count of shared zOTUs by treatment for tadpoles and metamorphs separately. We used the number of putative *Bd* inhibitor taxa shared between fathers and offspring per set, as the response variable, and treatment (normal transport, no transport, foster transport) as the independent variable. For the tadpoles we used a Poisson distribution and for metamorphs a negative binomial distribution.

All GLM and GLMM models were diagnosed using the *simulateResiduals* function in the DHARMa “package” version 0.4.7 ^14^. Additionally, we performed an Analysis of Deviance (Type II Wald chi square test) on our models to test for the significance of the independent variables using the function *gmmTMB:::Anova.glmmTMB* (“glmmTMB” package) for GLMMTMBs and the *Anova* function (“car” package, version 3.1-3, ^15^) for GLMs. Both Anova tests were followed by paired comparisons using the *emmeans* function in the “*emmeans*” package version 1.10.7 ^16^.

### Sample type composition

We found that the microbial community of all our samples was largely dominated by the phylum Proteobacteria (44 - 83 %), followed by the phyla Bacteriodota and Verrucomicrobiota in different proportions (Fig 2B; Supplementary material S4). The paternal samples were dominated by Proteobacteria (50.42%), Actinobacteriota (11.91 %), Bacteroidota (11.60%), Verrucomicrobiota (5.20%) and Firmicutes (5.02%). The maternal samples were dominated by the same phyla as the father, but in different proportions: Proteobacteria (50.42 %), Bacteroidota (13.20%), Actinobacteriota (13.20%), Verrucomicrobiota (4.66%) and Firmicutes (3.42%). For the dominant taxa of the tank samples, we found a slight difference in composition between substrate and water samples with the phyla Proteobacteria (substrate 47%; water 83 %), Actinobacteriota (substrate 11.78 %; water 1.14 %), Bacteroidota (substrate 11.47 %; water 9.54 %), Planctomycetota (substrate 5.30%; water 1.10 %) and Verrucomicrobiota (substrate 4.52 %; water 1.61 %). Offspring samples were dominated by Proteobacteria (w1 55.04%; w2 44.21%; tadpoles 63.31%; metamorphs 62.80%), Bacteroidota (w1 20.05%; w2 18.58%; tadpoles 12.82%; metamorphs 20.53%); Verrucomicrobiota (w1 10.22%; w2 22.02%; tadpoles 4.61%; metamorphs 4.22%), Actinobacteriota (w1 3.85%; metamorphs 3.64%), Bdellovibrionota (w1 3.06%; w2 2.21%), Planctomycetota (w2 4.12%; tadpoles 4.33%), and Chloroflexi (tadpoles 5.56%; metamorphs 1.89%). Across all samples, we found several taxa that have been previously identified as inhibitory for *Bd.* The genus *Aeromonas* was among the dominant taxa in samples of fathers (3.92 %), mothers (2.88 %), clutches (2.88 %), tadpoles (9.33 %), metamorphs (6.68 %) and substrate (2.72 %); the genus *Pseudomonas* was very abundant among fathers (5 %), mothers (4.48 %), clutches (4.48 %), metamorphs (7.01 %) and substrate (4.54 %); the genus *Shewanella* was among the dominant taxa in tadpole samples (7.11 %), and the genera *Pedobacter* (2.77 %) and *Flavobacterium* (2.31 %) were found to be very abundant in the late clutch samples. We did not find any taxa known to be inhibitory of *Bd* to be dominant in the water samples. See S4 tables for details.

**Table S1.** Summary of pairwise comparisons using an ANOSIM test on the calculated Bray-Curtis distances between sample type composition. We applied a Bonferroni correction to account for multiple comparisons. Significant values highlighted in bold and with an *.

| Pairs | anosimR | *p*-value | p-adj |
| --- | --- | --- | --- |
| Male.vs.Female | -0.03 | 0.990 | 1 |
| Male.vs.Substrate | 0.05 | 0.107 | 1 |
| Male.vs.Clutch_W1 | 0.28 | 0.002 | 0.056 |
| Male.vs.Water | 0.38 | 0.001 | **0.028\*** |
| Male.vs.Clutch_W2 | 0.48 | 0.001 | **0.028\*** |
| Male.vs.Tadpole | 0.67 | 0.001 | **0.028\*** |
| Male.vs.Metamorph | 0.29 | 0.001 | **0.028\*** |
| Female.vs.Substrate | 0.02 | 0.172 | 1 |
| Female.vs.Clutch_W1 | 0.27 | 0.001 | **0.028\*** |
| Female.vs.Water | 0.38 | 0.001 | **0.028\*** |
| Female.vs.Clutch_W2 | 0.47 | 0.001 | **0.028\*** |
| Female.vs.Tadpole | 0.64 | 0.001 | **0.028\*** |
| Female.vs.Metamorph | 0.26 | 0.001 | **0.028\*** |
| Substrate.vs.Clutch_W1 | 0.22 | 0.002 | 0.056 |
| Substrate.vs.Water | 0.38 | 0.001 | **0.028\*** |
| Substrate.vs.Clutch_W2 | 0.45 | 0.001 | **0.028\*** |
| Substrate.vs.Tadpole | 0.70 | 0.001 | **0.028\*** |
| Substrate.vs.Metamorph | 0.38 | 0.001 | **0.028\*** |
| Clutch_W1.vs.Water | 0.40 | 0.001 | **0.028\*** |
| Clutch_W1.vs.Clutch_W2 | 0.11 | 0.009 | 0.252 |
| Clutch_W1.vs.Tadpole | 0.77 | 0.001 | **0.028\*** |
| Clutch_W1.vs.Metamorph | 0.46 | 0.001 | **0.028\*** |
| Water.vs.Clutch_W2 | 0.53 | 0.001 | **0.028\*** |
| Water.vs.Tadpole | 0.80 | 0.001 | **0.028\*** |
| Water.vs.Metamorph | 0.58 | 0.001 | **0.028\*** |
| Clutch_W2.vs.Tadpole | 0.82 | 0.001 | **0.028\*** |
| Clutch_W2.vs.Metamorph | 0.59 | 0.001 | **0.028\*** |
| Tadpole.vs.Metamorph | 0.39 | 0.001 | **0.028\*** |

### Alpha diversity results

When looking at overall alpha-diversity, we found significant differences between sample types (Kruskal-Wallis, Chao1 X2 = 57.68, df = 7, *p* < 0.01; Shannon, X2 = 85.34, df = 7, *p* < 0.01; Inverse Simpson X2 = 102.14, df = 7, *p* < 0.01). Therefore, we explored pairwise comparisons between sample types (Table S2).

**Table S2.**
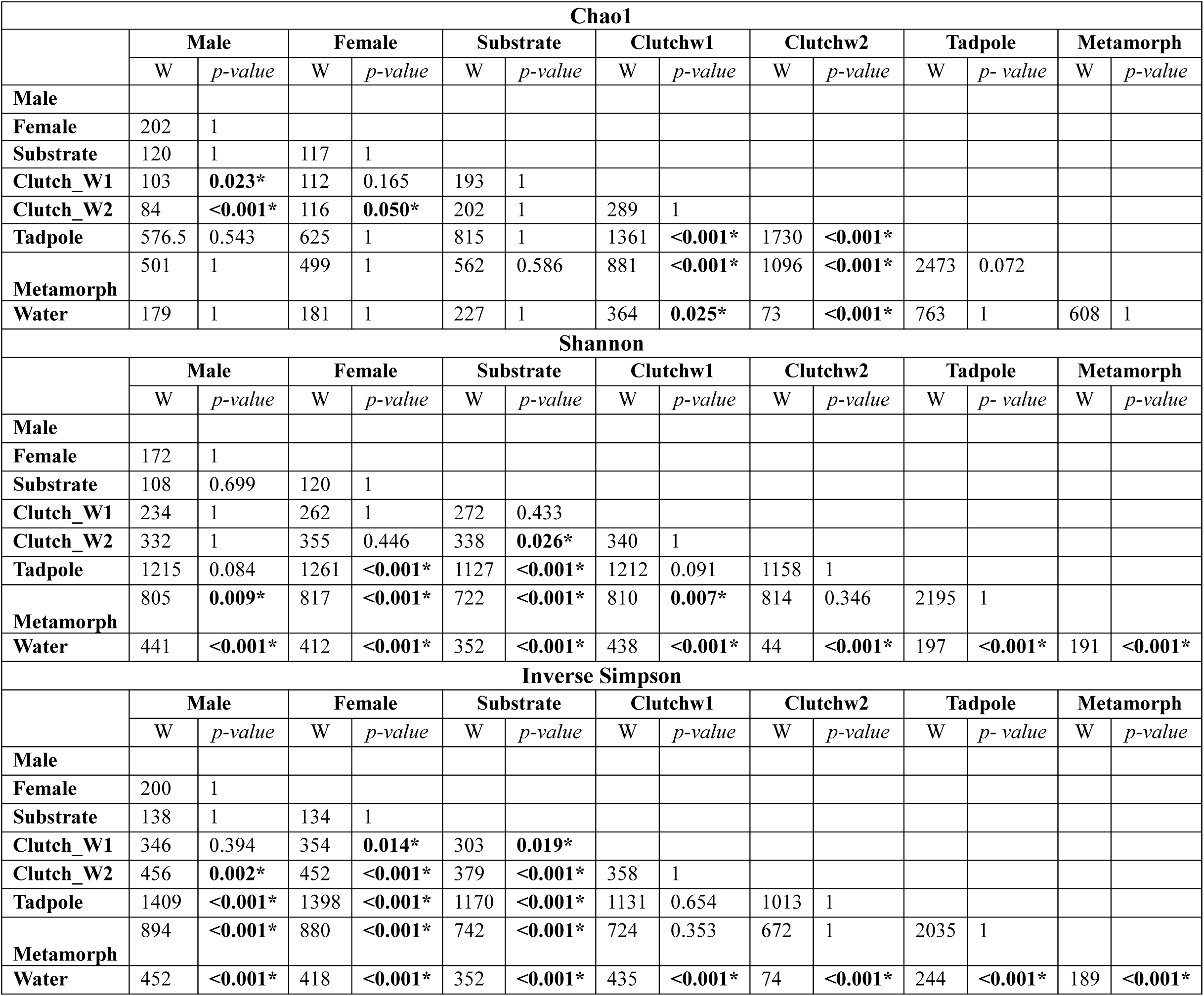
Pairwise comparisons of alpha-diversity indexes Chao1, Shannon and Inverse Simpson between sample types. Significant values highlighted in bold and with an *. W = Kruskal-Wallis estimate.

When investigating the effect of treatment on alpha-diversity in offspring samples (clutches w1 and w2, tadpoles, and metamorphs), only within tadpole samples for the Shannon Index we found statistically significant differences between treatments (Table S3). The pairwise comparisons revealed that tadpoles of the “no transport” treatment had a higher alpha diversity than tadpoles of the “foster transport” treatment (W = 4 243, *p =* 0.05, Table S4). However, the p-value was very close to 0.05, and because the other two alpha diversity measures did not reveal such a difference in tadpole samples across treatments, we assume that this difference is a statistical artefact, probably driven by two low outlier values in the “foster transport” treatment, and not a biologically relevant difference (Figure S3).

**Table S3.** Summary of tests (ANOVA and Kruskal-Wallis) of the effect of treatment on the alpha-diversity indexes found for offspring samples. Significant values highlighted in bold and with an *; df = degrees of freedom.

| Offspring sinks | Test type | Estimate | df | p-value |
| --- | --- | --- | --- | --- |
| *Clutch w1* |  |  |  |  |
| Chao1 | Anova | $F = 0.39$ | 2 | 0.683 |
| Shannon | Anova | $F = 0.13$ | 2 | 0.881 |
| Inverse Simpson | Kruskal-Wallis | $X^2 = 0.22$ | 2 | 0.894 |
| *Clutch w2* |  |  |  |  |
| Chao1 | Anova | $F = 1.14$ | 2 | 0.338 |
| Shannon | Anova | $F = 2.69$ | 2 | 0.090 |
| Inverse Simpson | Anova | $F = 3.43$ | 2 | 0.05 |
| *Tadpole* |  |  |  |  |
| Chao1 | Kruskal-Wallis | $X^2 = 0.97$ | 2 | 0.614 |
| Shannon | Kruskal-Wallis | $X^2 = 6.18$ | 2 | **0.045** |
| Inverse Simpson | Kruskal-Wallis | $X^2 = 4.14$ | 2 | 0.126 |
| *Metamorph* |  |  |  |  |
| Chao1 | Kruskal-Wallis | $X^2 = 4.55$ | 2 | 0.103 |
| Shannon | Kruskal-Wallis | $X^2 = 0.78$ | 2 | 0.678 |
| Inverse Simpson | Kruskal-Wallis | $X^2 = 0.10$ | 2 | 0.952 |

**Table S4.**
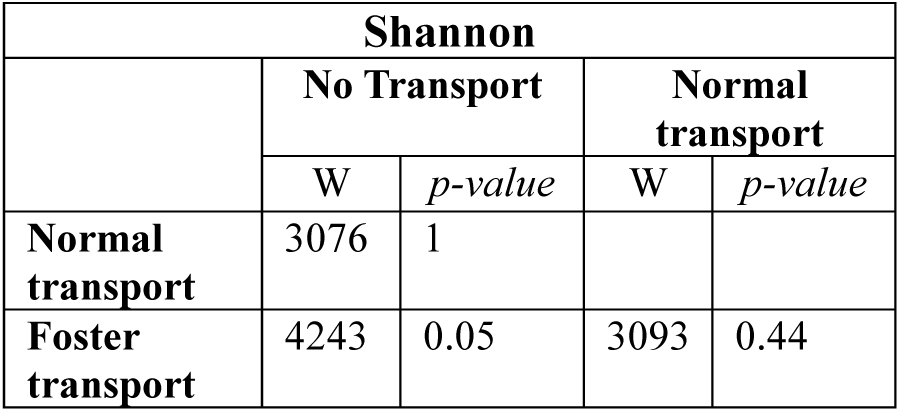
Pairwise comparisons of effect of treatment in tadpole samples for Shannon index. W = Kruskal-Wallis estimate.

**Table S5.** Sourcing analyses. Summary of percentages in contribution to sinks (offspring) from all sources tested.

| Treatment | Sink | Substrate<br>*Mean(SE)* | Mother<br>*Mean(SE)* | Biological<br>Father<br>*Mean(SE)* | Foster<br>Father<br>*Mean(SE)* | Water<br>*Mean(SE)* | Unknown<br>*Mean(SE)* | N |
| --- | --- | --- | --- | --- | --- | --- | --- | --- |
| Normal transport | *Clutch w1* | 9.75 (9.43) | 7.94 (5.82) | 26.43 (13.64) | - | 3.72 (2.67) | 52.15 (31.18) | 2 |
|  | *Clutch w2* | 5.63 (5.94) | 3.44 (2.63) | 3.59 (2.11) | - | 6.00 (4.85) | 81.35 (10.48) | 4 |
|  | *Tadpole* | 13.79 (12.96) | 5.66 (6.64) | 11.69 (9.16) | - | 11.34 (7.10) | 57.52 (11.41) | 8 |
|  | *Metamorph* | 12.47 (12.61) | 13.12 (12.37) | 8.01 (5.64) | - | 15.83 (18.38) | 50.58 (24.13) | 8 |
| No transport | *Clutch w1* | 6.03 (3.76) | 9.58 (8.28) | 6.46 (5.41) | - | 9.52 (12.74) | 68.40 (18.77) | 5 |
|  | *Clutch w2* | 3.62 (2.88) | 3.82 (4.02) | 4.35 (4.07) | - | 8.87 (7.78) | 79.35 (14.50) | 5 |
|  | *Tadpole* | 8.14 (9.43) | 10.10 (8.75) | 9.06 (9.53) | - | 13.95 (11.66) | 58.74 (10.37) | 22 |
|  | *Metamorph* | 11.24 (10.34) | 19.88 (13.73) | 6.80 (8.50) | - | 7.84 (8.97) | 54.24 (22.55) | 10 |
| Foster transport | *Clutch w1* | 9.48 (13.88) | 6.45 (6.54) | 7.58 (12.64) | 6.58 (9.16) | 8.66 (8.18) | 61.26 (25.33) | 4 |
|  | *Clutch w2* | 8.83 (10.95) | 5.43 (5.93) | 5.09 (4.02) | 4.07 (5.56) | 7.07 (12.39) | 69.51 (26.82) | 5 |
|  | *Tadpole* | 14.80 (17.43) | 26.93 (30.52) | 2.94 (2.65) | 14.02 (16.53) | 11.94 (9.97) | 29.38 (24.59) | 14 |
|  | *Metamorph* | 11.48 (12.10) | 18.95 (15.99) | 12.37 (8.34) | 12.61 (12.28) | 7.53 (8.73) | 37.06 (22.04) | 8 |
| Treatment | Developmental step | Contribution<br>*Mean (SE)* |  |  | Unknown Sources<br>*Mean (SE)* |  | N |  |
| Normal transport | *Embryonic development* | 29.24 (22.86) |  |  | 70.76 (22.86) |  | 7 |  |
|  | *Hatching* | 4.37 (3.50) |  |  | 95.63 (3.50) |  | 25 |  |
|  | *Metamorphosis* | 32.13 (20.87) |  |  | 67.87 (20.87) |  | 16 |  |
| No transport | *Embryonic development* | 56.60 (28.98) |  |  | 43.40 (28.98) |  | 4 |  |
|  | *Hatching* | 7.95 (12.27) |  |  | 92.05 (12.27) |  | 14 |  |
|  | *Metamorphosis* | 34.95 (16.01) |  |  | 65.05 (16.01) |  | 11 |  |
| Foster transport | *Embryonic development* | 33.54 (26.90) |  |  | 66.46 (26.90) |  | 9 |  |
|  | *Hatching* | 8.57 (9.39) |  |  | 91.43 (9.39) |  | 27 |  |
|  | *Metamorphosis* | 32.69 (17.93) |  |  | 67.31 (17.93) |  | 17 |  |

**Table S6.** ANOVA summary of Generalized Linear Mixed Model (glmmTMB) to determine differences in variables affecting contribution proportions. Significant *p*-values highlighted in bold.

| **Parameter** | **$X^2$** | **df** | ***p*-value** |
| --- | --- | --- | --- |
| *Treatment* | 0.47 | 2 | 0.789 |
| *Source* | 5.15 | 4 | 0.272 |
| *Sink* | 13.23 | 3 | **0.004** |
| *Treatment:Source* | 6.71 | 6 | 0.348 |
| *Treatment:Sink* | 3.94 | 6 | 0.685 |
| *Source:Sink* | 15.54 | 12 | 0.213 |
| *Treatment:Source:Sink* | 12.35 | 18 | 0.829 |

### Diagnostic plot for glmmTMB model testing differences in proportion of contribution of the known sources on the offspring

**Figure S2.**
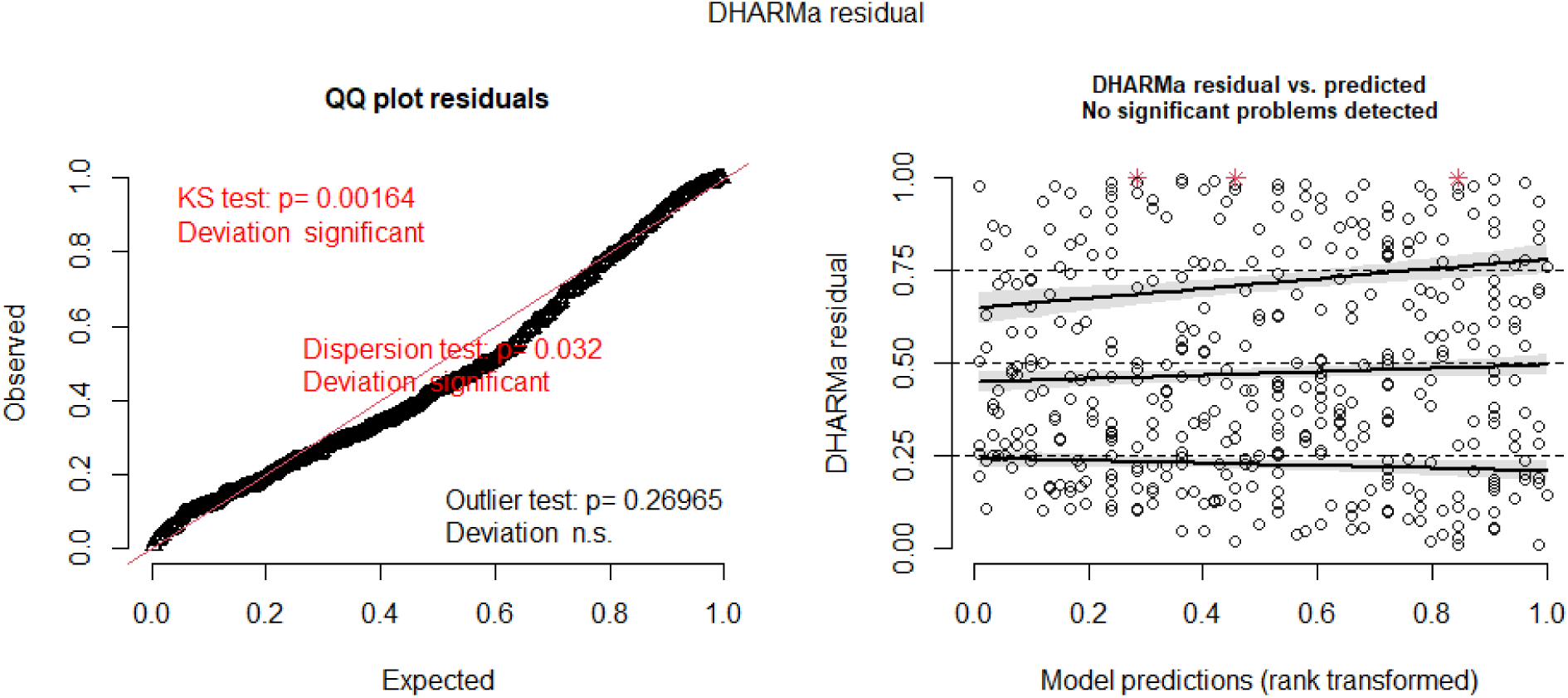
Plot of residual and predicted values using DHARMa package ^14^. We identified significant deviations from model assumptions (overdispersion: p = 0.016; uniformity: p = 0.014), however, this can be explained by the large sample size of our dataset and the sensitivity of the DHARMa test. Visual inspection of the QQ plot residuals suggest that the distribution of residuals is very close to the expected optimal values.

**Figure S3.**
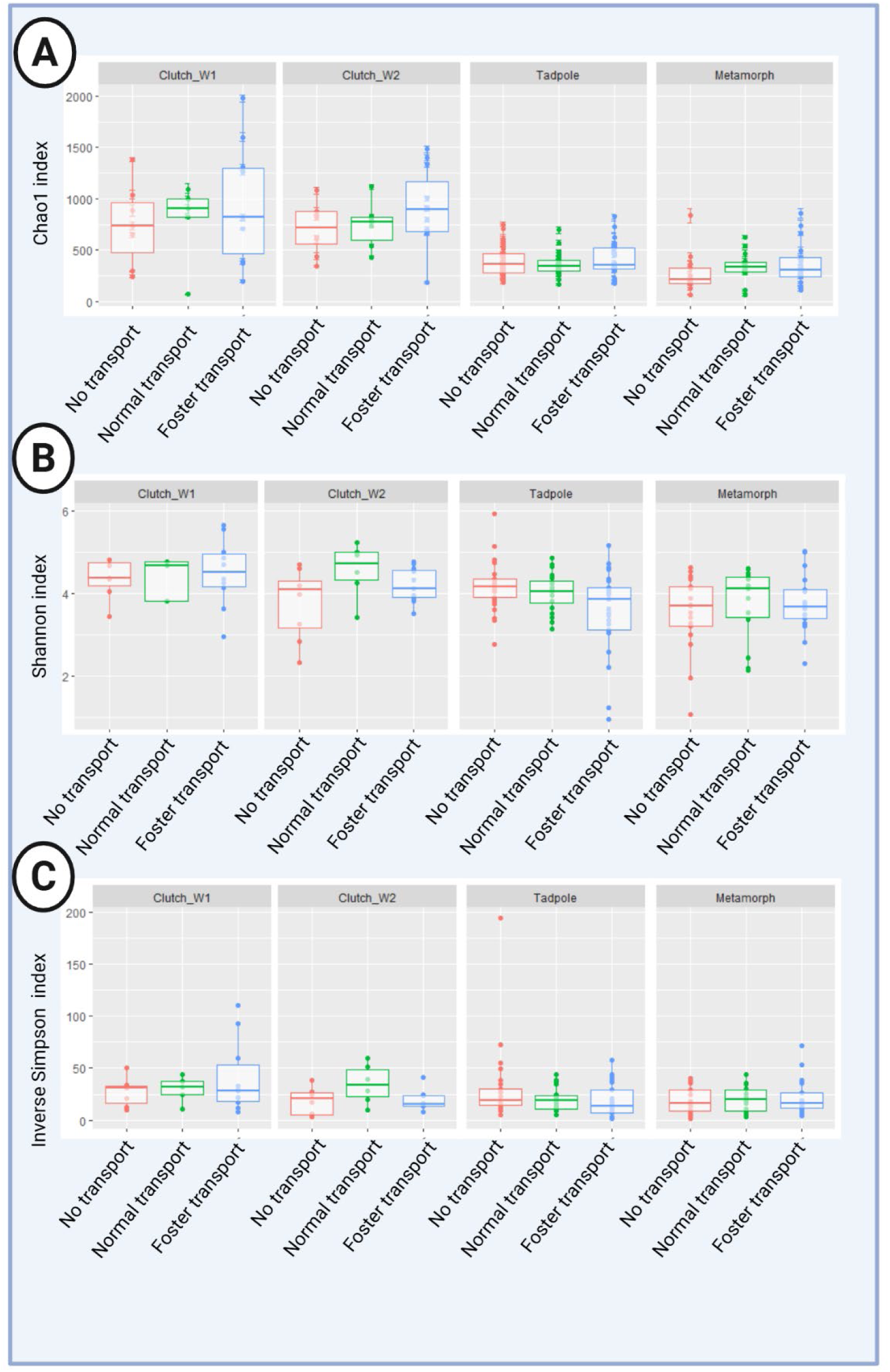
Boxplot of alpha diversity indexes of offspring samples. Alpha diversity measures A) Chao, B) Shannon, and C) Inverse Simpson Indexes are given for offspring samples per treatment. Created in BioRender. Angiolani Larrea, F. (2025) https://BioRender.com/2xtx2og

